# Genomic Underground: Unraveling NUMTs in Mole Voles

**DOI:** 10.1101/2023.12.30.573699

**Authors:** Dmitry Prokopov, Tigran Saluev, Svetlana Romanenko, Irina Bakloushinskaya, Alexander Graphodatsky

## Abstract

Nuclear mitochondrial DNA segments (NUMTs) are pervasive elements of eukaryotic genomes. This study focuses on *Ellobius talpinus* and *Ellobius lutescens*, for which we assembled full mitochondrial DNA sequences. Our study identified NUMTs encompassing approximately 0.0052% and 0.0086% of genome assembly length in *E. talpinus* and *E. lutescens*, respectively. These NUMTs collectively spanned a total length of 122,294 bp in *E. talpinus* and 194,875 bp in *E. lutescens*. Notably, the majority of NUMTs in both species were short, with lengths of less than 500 bp. In *E. talpinus*, the data indicated the presence of comparatively recent NUMT insertions. More than half of the NUMTs in each species are organized into clusters, primarily situated in intergenic regions or within introns. RNA genes are the most frequently occurring fragments within these NUMTs. Furthermore, our analysis identified LINE, SINE, and LTR retrotransposons within and flanking NUMT clusters. Our results demonstrate the intricate dynamics of NUMT integration and distribution in *Ellobius* species and provide insights into their genomic architecture and evolutionary history. This study contributes to the broader understanding of mitochondrial DNA contributions to nuclear genomes and underscores the complexity of distinguishing between mtDNA and nuclear DNA in genomic studies.

## Introduction

Mole voles (*Ellobius*, Arvicolinae, Cricetidae) are highly adapted to underground life social subterranean rodents that inhabit south-eastern Europe south of Western Siberia, Central Asia, and Mongolia^1–3^. Their genomic landscape, marked by unique features such as a distinct sex chromosome system^4–8^ and highly rearranged genomes with numerous inversions and translocations^9–11^, alludes to a rapid phase of genomic evolution within these subterranean rodents.

To date, the complete mitochondrial genomes of four *Ellobius* species have been sequenced: *Ellobius lutescens, Ellobius fuscocapillus, Ellobius talpinus*^12^, and *Ellobius tancrei*^13^, providing a foundation for understanding their mtDNA structure and evolutionary dynamics. Although their mitochondrial DNAs (mtDNAs) are typical of the Arvicolinae subfamily, they exhibit a unique feature of high GC content^14^. Recent studies^15,16^ have highlighted adaptive evolution in the mitochondrial genes of *Ellobius* species, particularly ATP8 and CYTB. These findings show varied selection pressures and link mtDNA evolution to rodent subterranean adaptations.

A notable aspect of genomic research is the study of nuclear mitochondrial DNA segments (NUMTs), which are fragments of mitochondrial DNA integrated into the nuclear genome. Discovered after sequencing the first human mitochondrial genome in 1983^17^, NUMTs were believed to traverse the nucleus through various mechanisms, such as homologous recombination^18^, non-homologous end joining^19^, microhomology-mediated end joining^20^, and retrotransposition^21^.

Although NUMTs are prevalent across eukaryotic genomes, the largest number of NUMTs has been described in mammals^20,22–29^. These repeated sequences can be utilized for various research purposes, such as population genetics^30^, past demographic event detection^31^, inference of hybridization events^32^, and biogeographic analysis^33^.

The presence of NUMTs poses difficulties in studying mitochondrial DNA (mtDNA), which may lead to the amplification of nuclear mtDNA copies. A recent study identified 195 NUMTs in the *E. talpinus* nuclear genome^34^. The high number of NUMTs highlights the challenges in differentiating mtDNA sequences and warns about the amplification of cryptic NUMTs in future studies on *E. talpinus*.

In this study, we conducted a thorough analysis of publicly available genomic data for *E. lutescens* and *E. talpinus*, using cutting-edge bioinformatics techniques. These techniques include mitochondrial DNA assembly, annotation, and NUMT detection. Additionally, we performed a comprehensive genome annotation to investigate the distribution, frequency, and structural patterns of NUMTs in these species.

## Materials and Methods

### Mitochondrial DNA assembly and annotation

Genomic reads were downloaded from the NCBI SRA database under accession numbers listed in Table 1.

**Table 1.** List of reads used in this study.

| Sample name | Species | SRA Accession | Tissue | Size (in Gb) | Study |
| --- | --- | --- | --- | --- | --- |
| ELUT 1 | *Ellobius lutescens* | SRR10542531 | Muscle | 4.4 | <sup>12</sup> |
| ELUT 2 |  | SRR3497433 | Liver | 35.7 | <sup>7</sup> |
| ELUT 3 |  | SRR3497460 | Liver | 20.1 | <sup>7</sup> |
| ETAL 1 | *Ellobius talpinus* | SRR10542532 | Muscle | 3.5 | <sup>12</sup> |
| ETAL 2 |  | SRR3497471 | Liver | 37.3 | <sup>7</sup> |

To ensure quality control, low-quality sequences, adapter sequences and PCR duplicates were removed using fastp 0.23.4^35^ with the following parameters: --detect_adapter_for_pe -D -- dup_calc_accuracy 6 -5 -3 -r -l 20. High-quality trimmed deduplicated reads were used for the mitogenome assembly. We used the assembler GetOrganelle v1.7.7.0^36^ with the option -F animal_mt to assemble mitogenomes. All available mitochondrial sequences of *Ellobius* from GenBank were used as the seed sequences. *de novo* assembly was manually checked and corrected. Protein-coding genes, non-coding regions, and RNA genes were annotated using MITOS2^37^. To detect mtDNA variants, the reads used for assembly were aligned against the newly assembled mitochondria using the BWA-MEM algorithm 0.7.17-r1188^38^. Variant calling was performed using the Ivar 1.4.2^39^.

### DNA Samples

Tissue samples of animals from a laboratory colony maintained in IDB RAS were used: four samples of *E. talpinus*, and four samples were used of *E. lutescens*. Tissue samples were deposited in the Large-Scale Research Facility “Collection of Wildlife tissues for genetic research” of the Core Centrum of the Koltzov Institute of Developmental Biology Russian Academy of Sciences (CWT IDB RAS), state registration number 3579666. The treatment of animals adhered to conventional international protocols in accordance with the guidelines for humane endpoints for animals used in biomedical research. The Ethics Committee for Animal Research of the Koltzov Institute of Developmental Biology RAS, which has approved all experimental protocols in compliance with the Regulations for Laboratory Practice in the Russian Federation (most recent protocol being № 37-25.06.2020), ensured that every possible measure was taken to minimize the animals’ suffering during capturing and sampling. Information on the origin founders of the colonies and sex of samples is presented in Table 2.

**Table 2.** Samples description

| № | Species | sex | Tissue | Location |
| --- | --- | --- | --- | --- |
| 1 | *Ellobius lutescens* | female | kidney | Armenia, near Vokhchabert settlement |
| 2 | *Ellobius lutescens* | female | kidney | Armenia, near Vokhchabert settlement |
| 3 | *Ellobius lutescens* | male | kidney | Armenia, near Atis settlement |
| 4 | *Ellobius lutescens* | male | kidney | Armenia, near Atis settlement |
| 5 | *Ellobius talpinus* | female | liver | Russia, Orenburg Oblast, near Alekseevka settlement |
| 6 | *Ellobius talpinus* | female | kidney | Russia, Saratov Oblast, near Khvalynsk |
| 7 | *Ellobius talpinus* | male | liver | Russia, Orenburg Oblast, near Alekseevka settlement |
| 8 | *Ellobius talpinus* | male | kidney | Russia, Saratov Oblast, near Khvalynsk |

DNA isolation was performed from tissues stored in ethanol using a Thermo Scientific™ GeneJET Genomic DNA Purification Kit (Thermo Fisher Scientific, USA) according to the manufacturer’s protocol.

### PCR

Unique specific primers were selected using Primer3 2.6.1 program^40^. The list of primer pairs is presented in Supplementary Table 1.

The PCR mixture contained 1**×** Taq buffer, 0.3 U Taq polymerase, 2 mM MgCl2, 200 µM dNTPs, 1 µM of each primer, and 20 ng of DNA.

To amplify the major non-coding region (NCR), located between mt-tRNA genes of phenylalanine and proline of *E. lutescens*, a protocol enabling the amplification of long DNA sequences (long PCR) was employed. Initial denaturation was performed at 94 °C for 3 min. Subsequently, the following cycling protocol was applied (35 cycles): 94 °C for 25 s, 68 °C for 10 s, and 68 °C for 2 min 30 s. The final elongation step was at 72 °C for 2 min and 30 s.

For the remaining amplification experiments, the program was as follows: initial denaturation at 94 °C for 3 min, followed by a cycling protocol (35 cycles) at 94 °C for 25 s, 68 °C for 10 s, and 68 °C for 50 s. The final elongation step was performed at 72 °C for 40 s.

Amplicons were visualized by electrophoresis on a 1.5% agarose gel using molecular markers S-8100 and S-8250 (BioLabMix).

### NUMTs detection in genome assembly

A similarity search was conducted on the nuclear and newly assembled mitochondrial genomes of two *Ellobius* species: male *E. lutescens* (ASM168507v1) and female *E. talpinus* (Etalpinus_0.1). Two mitochondrial genomes from each species were concatenated head-to-tail to avoid edge effects.

The search was performed using BLASTn version 2.15, with an e-value ≤ 5×10^-3^. NUMT sequences were concatenated into clusters if they followed each other on the scaffold.

### Genome annotation

The genomes of the mole voles from the NCBI Assembly database were not annotated; hence, we performed genomic annotation. For the identification and classification of repetitive sequences, we utilized RepeatModeler2 version 2.0.4, with the option -LTRStruct^41^ for *de novo* and *ab initio* repeat identification. Similar sequences of repeated elements were clustered using CD-HIT 4.8.1^42^. We used RepeatMasker version 4.1.5^43^ with the parameters -e rmblast -s -no_is to identify repetitive sequences in the whole-genome sequences. The coordinates of the repeats detected were subsequently intersected with the spacer sequences present in the NUMT clusters, as well as the flanking regions of these clusters, with lengths of 1,000 base pairs (bp).

Protein-coding gene structures were predicted *ab initio* using Augustus 3.4.0^44^ with the options -- strand=both --noInFrameStop=false --protein=on --codingseq=on --introns=on --start=on --stop=on --cds=on --singlestrand=false --UTR=off –genemodel=complete.

Manipulation with output tables and genomic intervals was performed using bedtools 2.31.1^45^ and custom scripts available at https://github.com/Saluev/ellobius.

## Results

### Assembled mtDNA

The GetOrganelle pipeline successfully generated completely circularized mitochondrial DNA sequences for all available NCBI SRA sample reads of *E. lutescens* and *E. talpinus*.

The assembled mitochondrial genomes were similar to those previously reported by Bondareva et al.^12^, although there were significant structural differences in *E. lutescens*. The newly assembled *E. lutescens* mitogenomes were considerably longer than those assembled in the study by Bondareva et al.^12^, with nucleotide differences ranging from 1,516 to 1,639 bp. The presence of light-strand replication origin (O_L_) and duplications in the control region, consisting of 2-3 duplicated origin for H-strand DNA replication (O_H_) sequences that had not been previously assembled, accounts for the difference in our assembly compared with previous studies.

To assess the copy number and size of the control region for both species, we performed PCR on the flanking regions and the internal segment of the O_H_. The observed amplicon length for *E. talpinus* perfectly matched the expected size determined by gel electrophoresis. However, for *E. lutescens,* the O_H_ amplicon was 2-3 times smaller than expected, and the copy number of this region was reduced by 2-3 times, suggesting assembly artifacts. Molecular validation of the O_H_ length allowed for manual trimming of the sequence, retaining only a single copy of the O_H_ sequence.

The assembly process was conducted for both ELUT3 and ELUT2 samples, originating from the same parental individuals^7^. This analysis revealed that ELUT2 and ELUT3 had identical maternal mitochondrial DNA, reflecting their shared maternal lineages. The characteristics of the assembled mitogenomes are listed in Table 3.

**Table 3.** Characteristics of the mitochondrial DNA of E. lutescens (ELUT) and E. talpinus (ETAL) from the NCBI database (coloured in gray) and this article.

| Name | Accession number | mtDNA length | A% | T% | G% | C% | GC-content | Control region |  |
| --- | --- | --- | --- | --- | --- | --- | --- | --- | --- |
|  |  |  |  |  |  |  |  | size | GC-content |
| ELUT NCBI | MT483992.1 | 16540 | 31.1 | 25.9 | 14.3 | 28.4 | 42.8 | 1172 | 41.5 |
| ELUT 1 |  | 16843 | 31.2 | 25.8 | 14.4 | 28.6 | 43 | 1169 | 41.4 |
| ELUT 2 and ELUT3 |  | 16853 | 31.1 | 25.8 | 14.5 | 29 | 43.1 | 1169 | 41.5 |
| ETAL NCBI | NC_054160.1 | 16367 | 32.4 | 26.3 | 13.6 | 27.7 | 41.3 | 976 | 39.8 |
| ETAL 1 |  | 16367 | 32.4 | 26.3 | 13.6 | 27.7 | 41.3 | 976 | 39.8 |
| ETAL 2 |  | 16361 | 32.8 | 26.5 | 13.3 | 27.4 | 40.7 | 969 | 38.9 |

### NUMTs identification

Bioinformatics analysis of the mitochondrial genome in the nuclear genomes of *E. lutescens* and *E. talpinus* revealed the presence of 167 and 463 fragments, respectively. The total length of NUMTs in the *E. talpinus* and *E. lutescens* genomes was 122 294 and 194 875 bp, respectively, which accounted for 0.0052% and 0.0086% of the genome assembly length, or 0.0041% and 0.0068% of the nuclear genome, respectively, based on previous genome amount estimations^46^. The majority of NUMTs exhibited a length of less than 500 bp (mean: 732 bp, median: 471 bp in *E. lutescens* and mean: 421 bp, median: 244 bp in *E. talpinus*), as depicted in Supplementary Figure 1. The degree of sequence identity between mtDNA and NUMT exhibits considerable variation, encompassing values spanning from 71.5% to 100%. The most frequently observed identity was registered at 80-84% (mean: 87%, median: 85%) in *Ellobius lutescens*, while 90-94% (mean: 90%, median: 90%) was recorded in *Ellobius talpinus* (Supplementary Figure 2).

The most frequently occurring fragments of the mtDNA sequence in the nuclear genome were RNA-encoding sequences (263 and 447 genes in NUMTs for *E. talpinus* and *E. lutescens,* respectively) and protein-coding genes (153 and 372). The lowest number of mtDNA fragments found in the nuclear genome was found in non-coding regions (Figure 1, Supplementary Figure 3). The most frequent genes in NUMTs were rrnL (40 fragments in *E. lutescens* and 80 in *E. talpinus*) and control regions (33 and 80 fragments, respectively).

**Figure 1:**
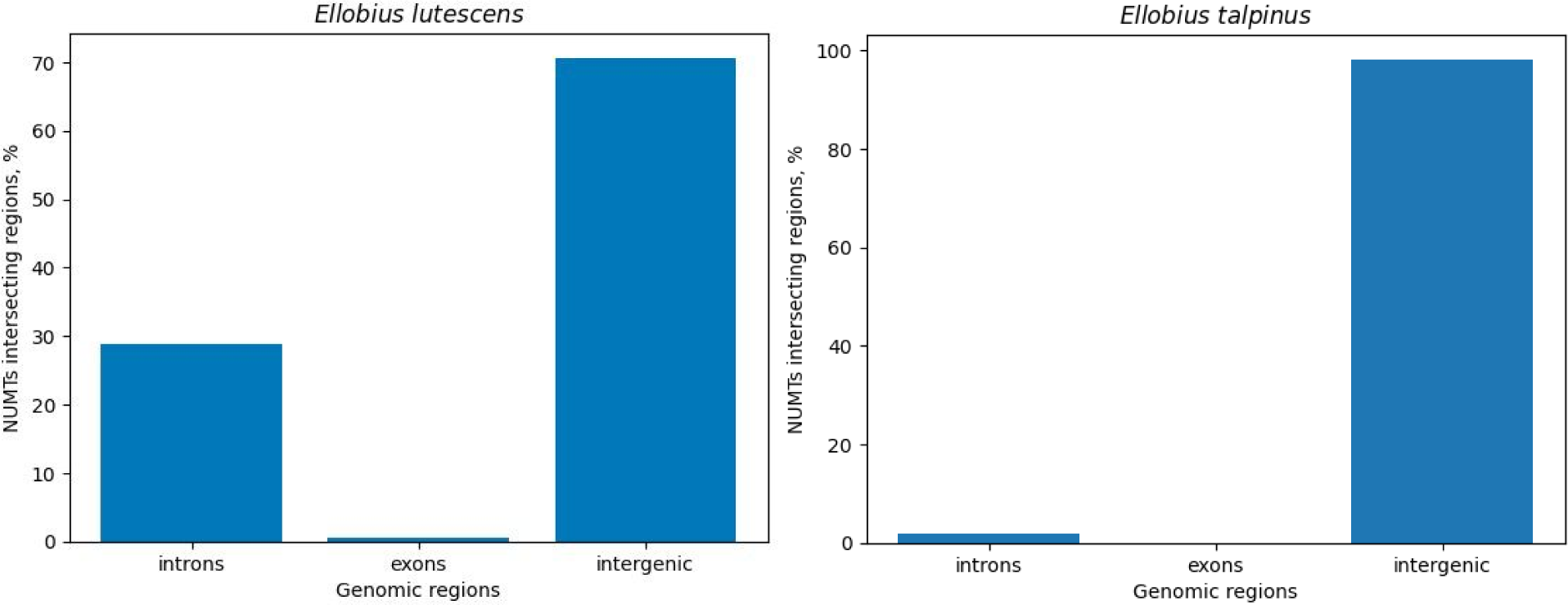
Visual representation of NUMT alignments on the mitochondrial DNA map of Ellobius lutescens and Ellobius talpinus. The arrangement of the sequences is demonstrated using arrows, with the direction of the sequence indicated. The colour scheme, which range from yellow to red, conveys the degree of identity.

51.5% and 45.6% of NUMTs in *E. lutescens* and *E. talpinus* organized in clusters (86 and 211 clusters in the genome, respectively), containing homologous mtDNA sequences interrupted by spacer sequences.

## Genomic context of NUMTs

Analysis of NUMTs in relation to genomic annotations revealed that the majority of NUMTs were located in intergenic regions of the genome, with a notable proportion of NUMTs detected in gene regions (Figure 2). Within these gene regions, introns are the most common location for NUMTs, followed by exons. In the genome of *E. lutescens*, only one NUMT was identified within the coding region of a gene, specifically the second exon of the PDSS2 truncated pseudogene.

**Figure 2:**
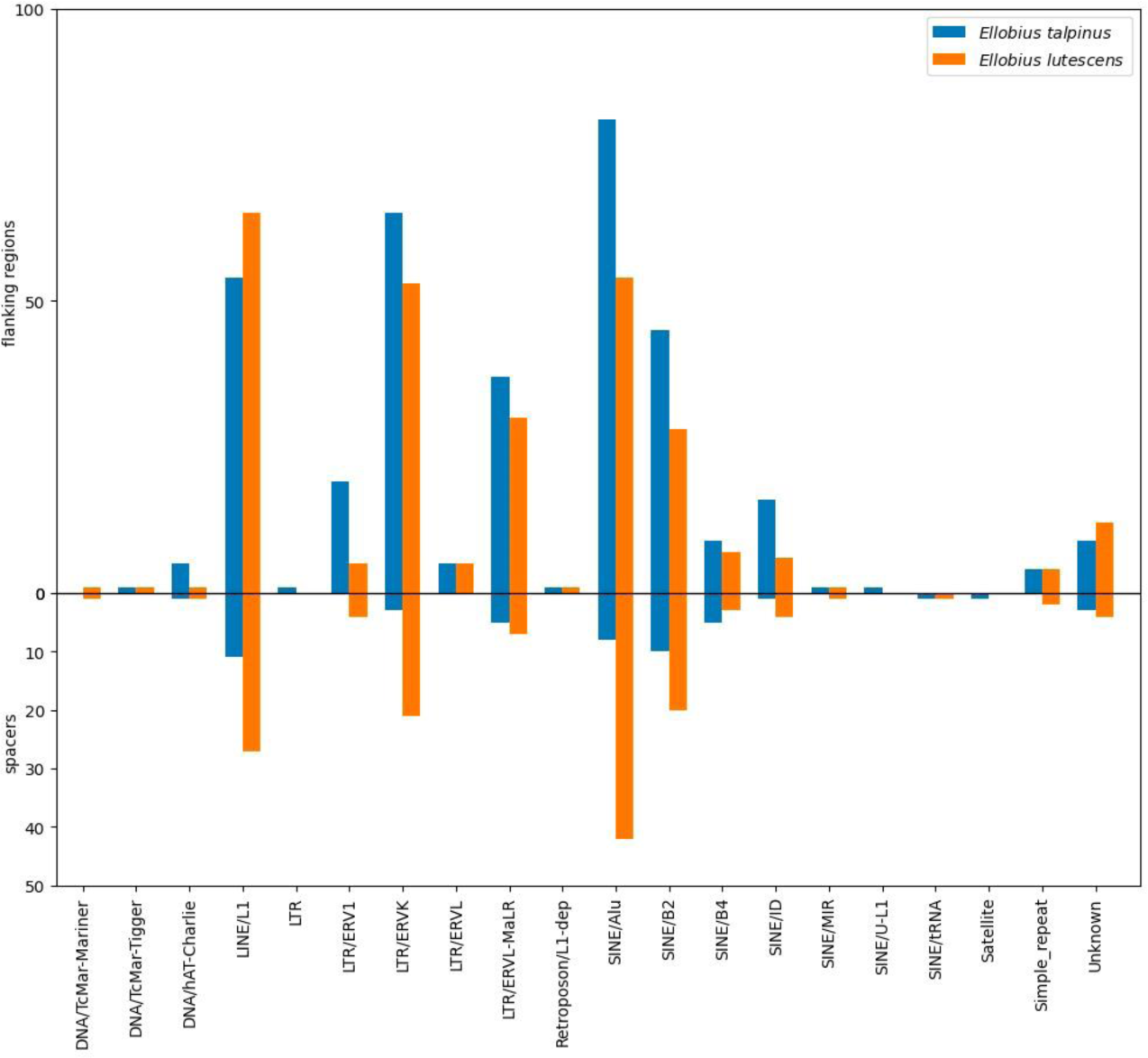
Location of NUMTs in the genomic regions of two mole vole species.

We investigated the occurrence of repetitive sequences both within and around NUMT clusters. (Figure 3).

**Figure 3:**
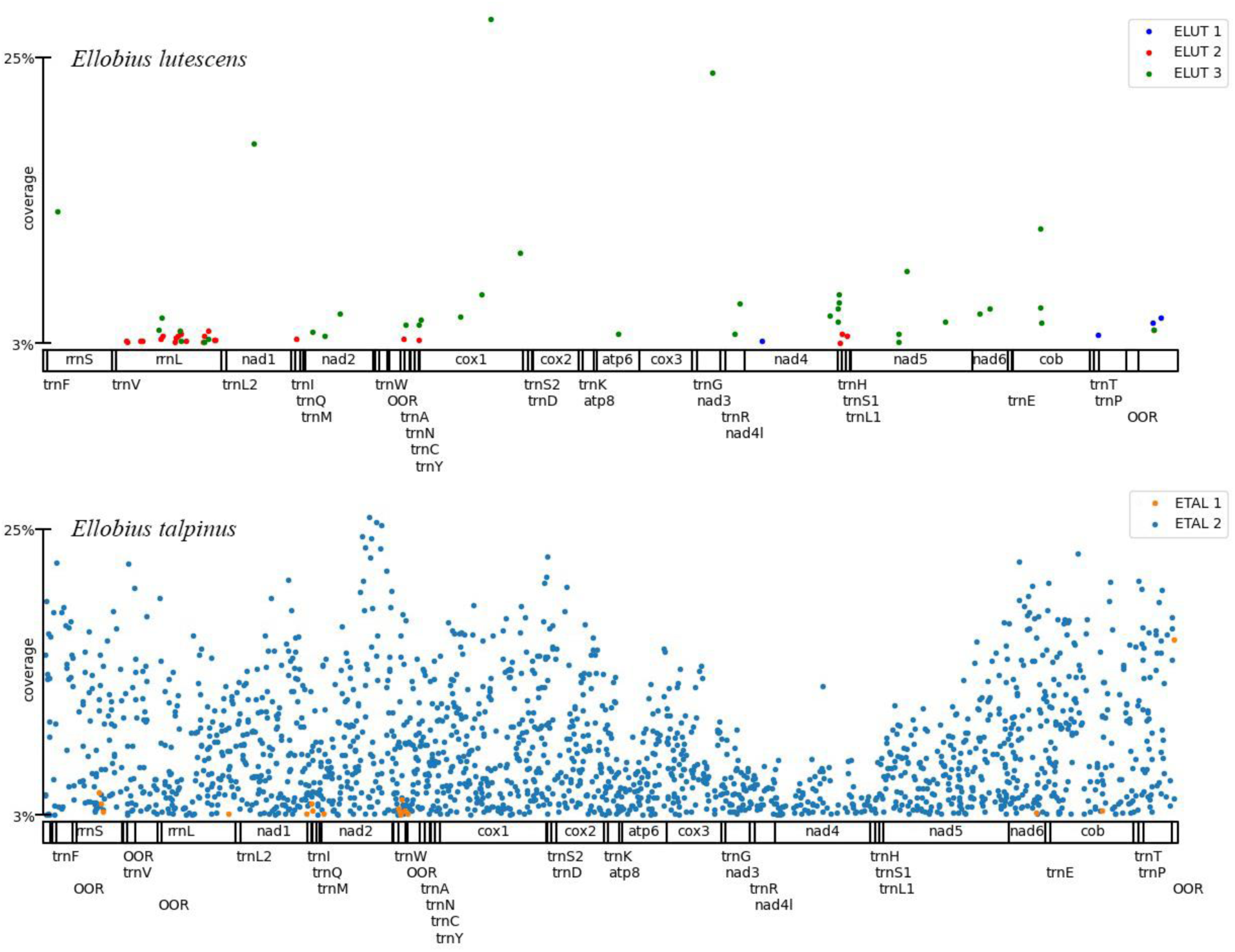
Content of repeated elements in the internal and flanking regions of NUMT clusters.

In *E. lutescens*, 166 repetitive sequences were found within the NUMT clusters, with LTRs (22.3%), SINEs (43.4%, mainly SINE/Alu), and LINEs (23.5%, all LINE/L1). The flanking regions contained 277 sequences, showing a higher prevalence of LTRs (34.7%, mainly LTR/ERVK) and SINEs (33.6%).

In *E. talpinus,* NUMT clusters contained 82 repeats, with a composition of LTRs (29.3%, mostly LTR/ERVL-MaLR and LTR/ERVK), SINEs (54.9%, predominantly SINE/Alu and SINE/B2), and LINEs (17.1%, all LINE/L1). The flanking areas had 322 sequences, with a larger proportion of LTRs (42.5%, mainly LTR/ERVK) and SINEs (43.8%).

### Nucleotide variants and coverage analysis

Several mitochondrial single-nucleotide variants (SNV) were identified in *E. lutescens* samples, of which five possessed a considerable degree of coverage (Figure 4).

**Figure 4:**
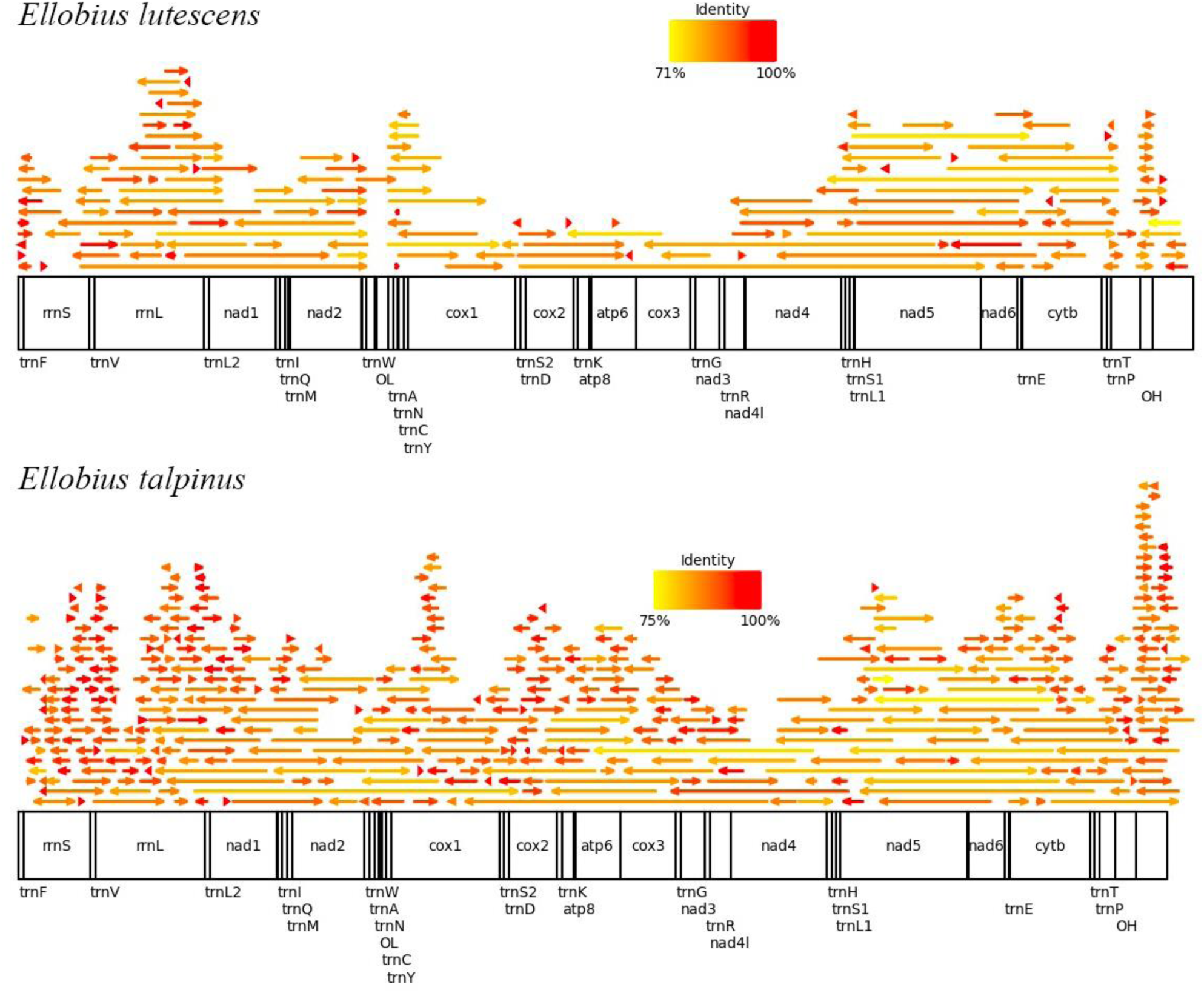
SNV composition in mitochondrial genomes of Ellobius lutescens and Ellobius talpinus.

An unusual observation was made regarding the *E. talpinus* samples. Specifically, in one sample (ETAL 2), a large number of nucleotide variants were detected with high coverage, which was similar to the distribution pattern of the NUMT clusters (Figure 1). Conversely, for ETAL 1, only a small number of nucleotide variants were observed, and these were mostly of low coverage (Figure 4).

## Discussion

Our study provides an in-depth look at the distinctive patterns of mitochondrial DNA in the nuclear genomes of *E. lutescens* and *E. talpinus*. A detailed exploration of NUMTs uncovers genomic complexities, offering insights into the dynamic interaction between mitochondrial and nuclear genomes. Furthermore, our analysis extends beyond mere identification, as we uncovered the composition, organization, and genomic context of NUMTs, offering novel insights into the evolutionary dynamics of these rodent species.

### Mitochondrial Genome Assembly Challenges: Lessons from *Ellobius lutescens*

To comprehensively study the NUMT genome composition, it is necessary to obtain *E. lutescens* and *E. talpinus* complete sequence mtDNA. The GetOrganelle pipeline enabled the successful assembly of circular mitochondrial DNA in both species. Previously, the mitochondrial genome of *E. lutescens* (MT483992.1) had two gaps of unknown nucleotide composition, with a total of 39 nucleotides. The new genomic assembly fills these gaps, resulting in the first complete mitochondrial assembly of this species.

However, the new assembly included extended tandem duplications in the control region (O_H_ sequence), which were repeated two-three times. Analysis of the duplicated sequence length using PCR suggested that this duplication was an artifact of the assembly process. It is probable that this duplication occurred because of the use of short reads and the De Bruin graph for sequence assembly^47^. Although GetOrganelle has demonstrated the highest accuracy among all published plastid assembly pipelines in benchmarks^48–50^, this observation underscores the importance of the molecular validation of *in silico* results.

### NUMT Dynamics Unveiled: Contrasting Abundance, Length, and Composition in *Ellobius lutescens* and *Ellobius talpinus*

Our analysis of NUMTs in *E. lutescens* and *E. talpinus* revealed noteworthy differences in abundance, total length, and composition. Specifically, *E. talpinus* displayed a higher number and total length of NUMTs, indicating greater propensity for NUMT integration. This disparity may be attributed to variations in the mechanisms and rates of NUMT transfer between mitochondrial and nuclear genomes. Previously, high NUMT polymorphisms in the genomes, in terms of both copy number and total length, were also observed in some closely related species of fig wasps^51^, fruit flies^52^, bats^27^, marsupial shrews^53^ and primates^25^.

The structuring of NUMTs into clusters with homologous mtDNA sequences interspersed by spacer sequences suggests multiple rounds of insertion or duplication, which may have been facilitated by non-homologous end-joining, replication-dependent DNA double-strand break repair and/or retrotransposition^21,54^. The presence of NUMT clusters has also been described in some plants^55^ and yeast^56^.

The predominance of short NUMTs could also be attributed to integration mechanism into the nuclear genome. During the transfer of mitochondrial DNA to the nuclear genome, longer fragments may be more prone to degradation or truncation. Additionally, the breakage and repair processes that facilitate the integration of mtDNA into the nuclear genome may favor shorter segments because of their easier incorporation and less disruptive impact on nuclear genome stability.

Analysis of the composition of RNA-encoded sequences in NUMTs revealed that they were the most prevalent fragments in both genomes. This led us to propose a hypothesis that the higher concentration of RNA genes, including tRNAs and rRNAs, compared to protein-coding genes, increases the likelihood of integration into the nuclear genome. The abundance of these RNA molecules in cytoplasm^57,58^ might lead to preferential transfer or integration of these sequences into the nuclear genome because of their availability.

The higher level of identity in *E. talpinus* NUMTs suggests a more recent transfer of these sequences to the nuclear genome. The high identity implies less time for mutations to accumulate in these NUMTs since their integration, suggesting a dynamic and ongoing process of mitochondrial DNA incorporation into the nuclear genome in *E. talpinus*.

### Decoding the NUMT Landscape: Genomic Signatures and Repetitive Element Associations

Examination of the genomic context of NUMTs provided noteworthy findings. NUMTs are predominantly located within the intergenic regions and are enriched in introns, a characteristic feature of most studied NUMTs^20,23,25,59,60^. This distribution suggests that NUMTs in these regions face less selective pressure than those within coding regions. The presence of NUMTs within introns may be due to lower functional constraints in these regions, allowing for their integration without significantly disrupting gene function.

Additionally, the presence of repeated elements, particularly LINE, SINE, and LTR, in the flanking regions of NUMTs, which has also been observed in various species^21,23,25^, suggests their possible involvement in NUMT insertion and mobilization.

### Deciphering SNV Complexity: Insights and Challenges in Mitochondrial Genomic Analysis

Mitochondrial single nucleotide variant (SNV) analysis in *E. lutescens* and *E. talpinus* samples revealed intriguing patterns. In *E. lutescens*, the detection of several SNVs, with four exhibiting high coverage, suggests subtle heteroplasmy. However, distinguishing mitochondrial variations from nuclear mitochondrial DNA poses challenges, necessitating advanced sequencing approaches, especially long-read data.

Caution is crucial when interpreting SNV data in *E. talpinus* because of the prevalence of NUMT sequences. Significant SNV coverage in ETAL 2 (Fugure 4) suggests a NUMT association, supported by distribution patterns resembling NUMT clusters (Figure 2). This highlights the challenges in accurate mitochondrial DNA analysis and underscores the need for careful consideration in species with high NUMT prevalence. A notable absence of SNVs from the NUMTs was observed in ETAL 1 (Table 1). Plausible reasons include low coverage and muscle tissue bias towards heteroplasmic mitochondrial variants owing to its higher mtDNA content. Choosing the muscle tissue for barcoding may enhance reliability by minimizing NUMT-related amplification bias. NUMT co-amplification is known to occur in species that have experienced a recent NUMT insertion and poses a major challenge to the interpretation of DNA barcoding results^61–71^. A recent study revealed the co-amplification of NUMT-derived sequences within a control region fragments in *E. talpinus*, which could result in erroneous outcomes in population analysis^34^. Therefore, it is essential to account for NUMTs in molecular studies and employ additional sequencing techniques and methodologies to ensure the fidelity of species barcoding.

## Conclusion

In this study of *Ellobius lutescens* and *Ellobius talpinus*, we systematically characterized NUMTs, revealing distinct patterns in their abundance, length, and genomic distribution. Our analysis indicates significant differences between the two species, with *E. talpinus* exhibiting more recent NUMT insertions. The prevalence of short NUMTs, often clustered and predominantly located in non-coding regions, suggests complex integration mechanisms. The identification of LINE, SINE, and LTR elements near NUMT clusters implies their role in NUMT mobilization. Our work highlights the challenges in distinguishing mitochondrial DNA from NUMTs in genomic studies, underscoring the need for careful data interpretation and advanced sequencing approaches. This research contributes to the understanding of NUMT dynamics in mole voles and their potential impact on genomic evolution.

## Contributions

D.P.: conceptualization; D.P.: methodology; T.S.: software, D.P. and S.R.: validation, D.P. and T.S.: formal analysis; D.P., I.B., T.S., and S.R.: investigation; A.G.: resources, D.P. and T.S.: data curation; D.P.: writing – original draft preparation, D.P., I.B., T.S., S.R., and A.G.: writing – review and editing; D.P., and T.S.: visualization; D.P.: supervision; S.R., and A.G.: project administration; A.G.: funding acquisition. All authors reviewed and approved the final manuscript.

## Corresponding author

Correspondence to Dmitry Prokopov.

## Supplementary files

**Genomic Underground: Unraveling NUMTs in Mole Voles**

Dmitry Prokopov, Tigran Saluev, Svetlana Romanenko, Irina Bakloushinskaya, Alexander Graphodatsky

**Supplementary Figure 1.**
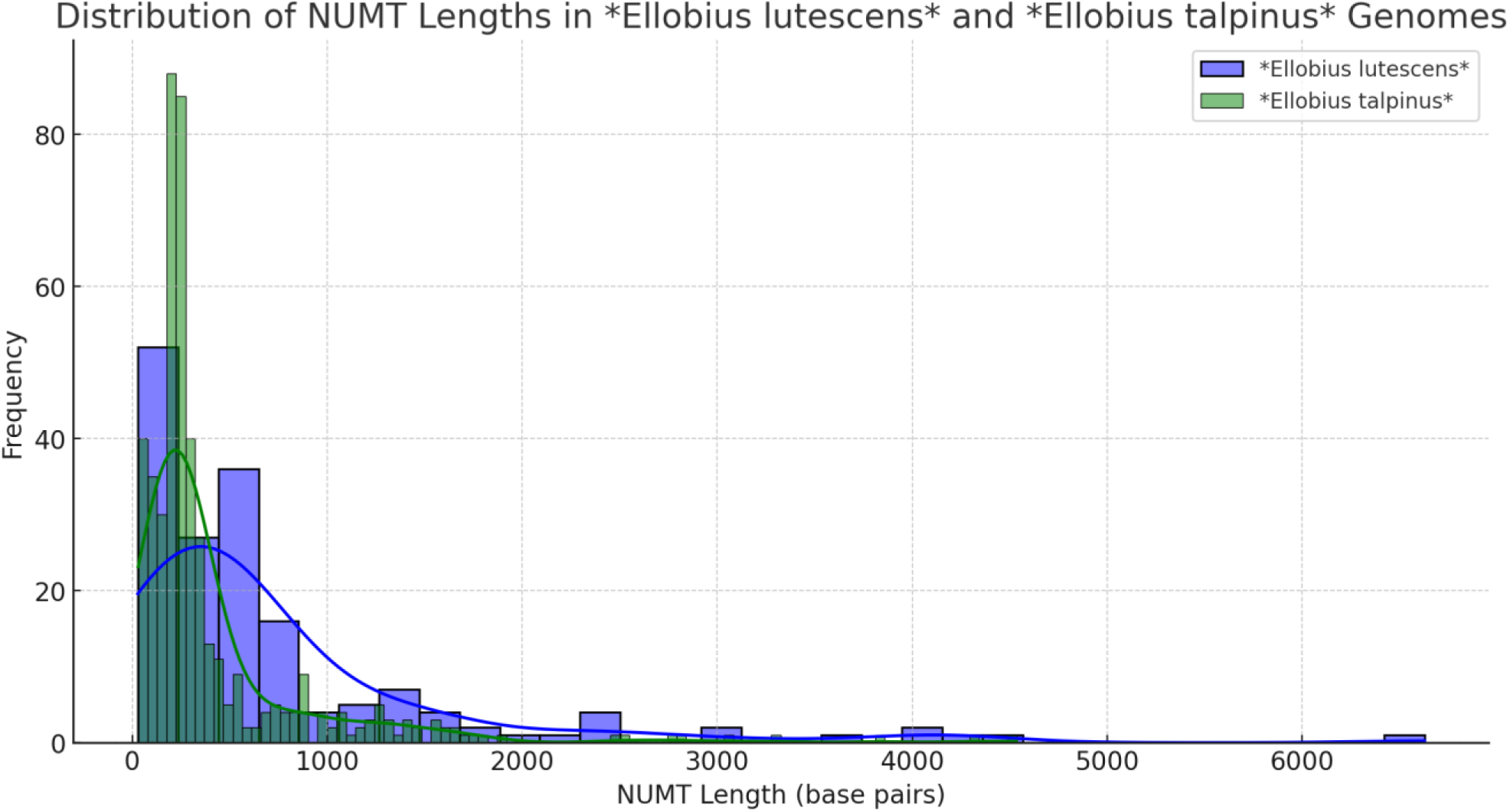
Distribution of NUMT lengths in the genomes of Ellobius lutescens (in blue) and Ellobius talpinus (in green). The x-axis represents the length of NUMTs in base pairs, and the y-axis shows the frequency of each length range.

**Supplementary Figure 2.**
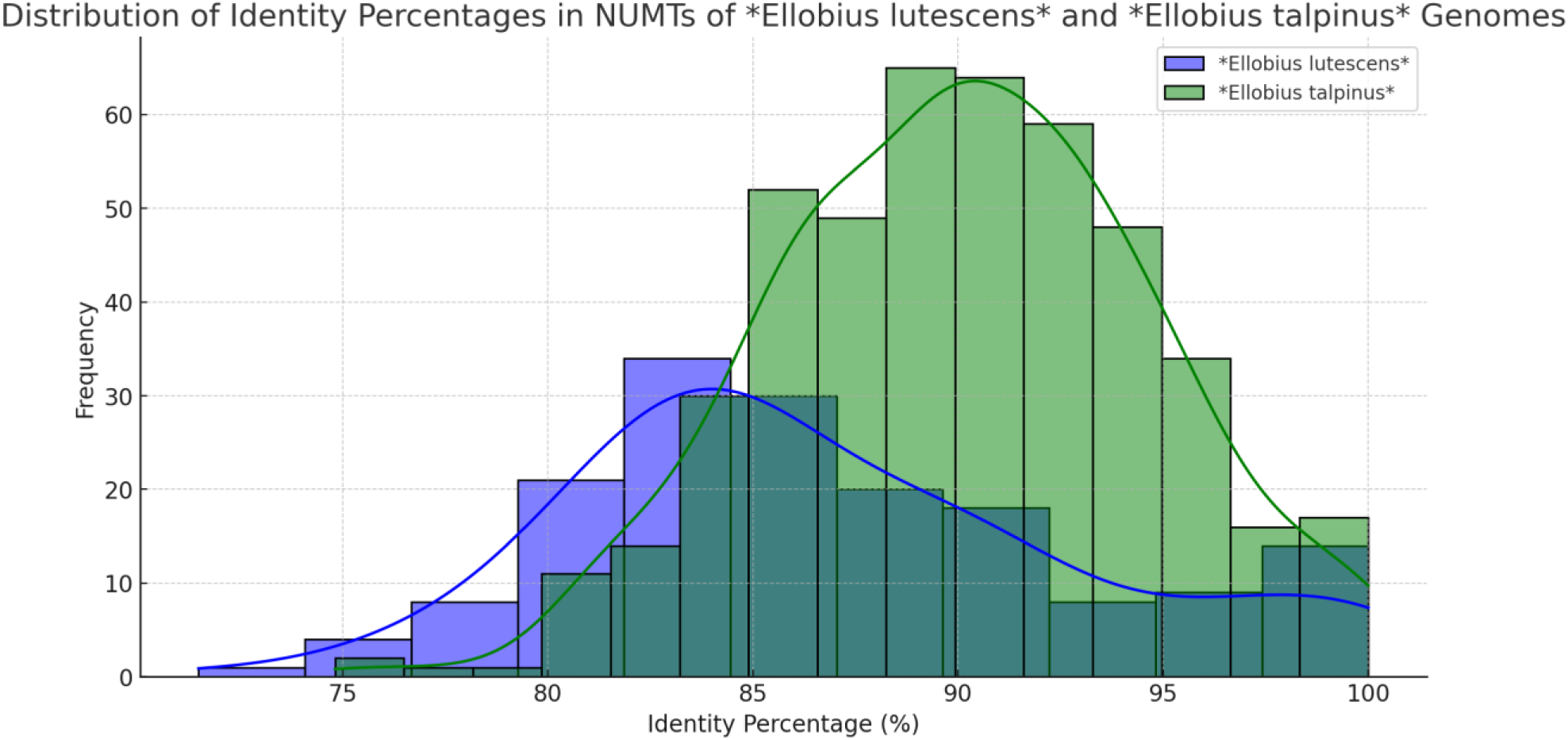
Distribution of identity percentages for NUMTs in the genomes of Ellobius lutescens (in blue) and Ellobius talpinus (in green). The x-axis represents the percentage of identity, and the y-axis shows the frequency of each identity range.

**Supplementary Figure 3.**
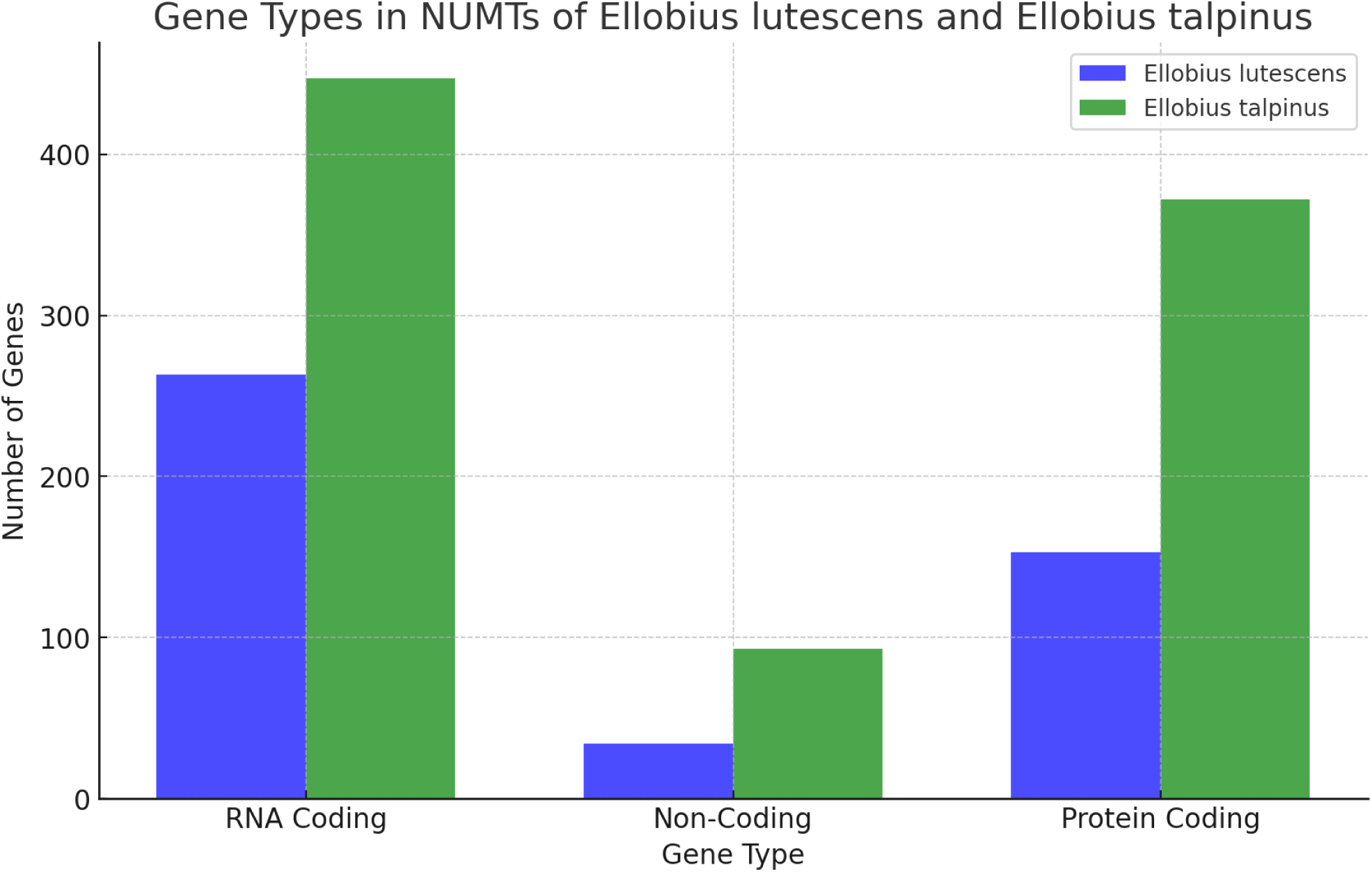
The types of genes present in NUMTs of Ellobius lutescens and Ellobius talpinus. The chart shows the number of RNA coding genes (including tRNA and rRNA), non-coding genes (OL and control region), and protein-coding genes in NUMTs for each species.

**Supplementary Table 1.**
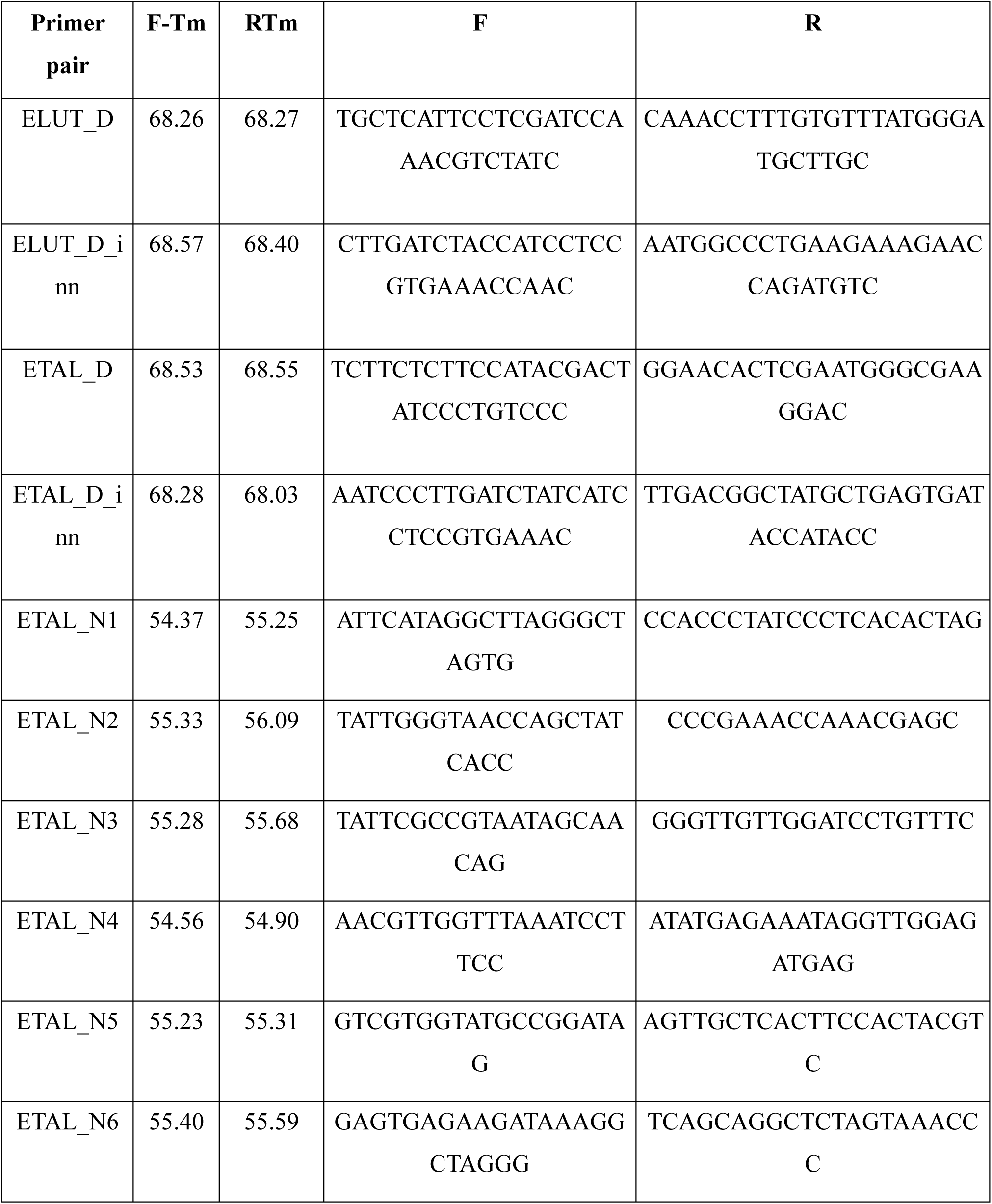
List of primer pairs

